# Oxytocin receptor is not required for social attachment in prairie voles

**DOI:** 10.1101/2022.07.22.501192

**Authors:** Kristen M. Berendzen, Ruchira Sharma, Maricruz Alvarado Mandujano, YiChao Wei, Forrest D. Rogers, Trenton C. Simmons, Adele H.M. Seelke, Jessica M. Bond, Rose D. Larios, Michael Sherman, Srinivas Parthasarathy, Isidero Espineda, Joseph R. Knoedler, Annaliese Beery, Karen L. Bales, Nirao M. Shah, Devanand S. Manoli

**Affiliations:** Department of Psychiatry and Behavioral Sciences, University of California, San Francisco; San Francisco, CA 95158, USA; Center for Integrative Neuroscience, University of California, San Francisco; San Francisco, CA 95158, USA; Weill Institute for Neurosciences, University of California, San Francisco; San Francisco, CA 95158, USA; Kavli Institute for Fundamental Neuroscience, University of California, San Francisco; San Francisco, CA 95158, USA; Department of Psychology, Physiology, and Behavior, University of California, Davis; Davis, CA 95616, USA; Neurosciences Graduate Program, University of California, San Francisco; San Francisco, CA 95158, USA; Department of Neurobiology, Physiology, and Behavior, University of California, Davis; Davis, CA 95616, USA; Department of Psychiatry and Behavioral Sciences, Stanford University; Stanford, CA 94305, USA; Department of Neurobiology, Stanford University; Stanford, CA 94305, USA; Department of Integrative Biology, University of California, Berkeley; Berkeley, CA 94720, USA

## Abstract

Prairie voles are among a small group of mammals that display long-term social attachment between mating partners. Many pharmacological studies show that signaling via the oxytocin receptor (OxtR) is critical for the display of social monogamy in these animals. We used CRISPR-mutagenesis to independently generate three different OxtR null mutant prairie vole lines. OxtR mutants displayed social attachment such that males and females showed a behavioral preference for their mating partners over a stranger of the opposite sex when assayed using different paradigms. Mothers lacking OxtR delivered viable pups, and parents displayed care of their young and raised them to the weanling stage. Together, our studies unexpectedly reveal that OxtR-mediated signaling is genetically dispensable for social attachment, parturition, and parental behavior.

## Introduction

Strong, specific, and sustained relationships between mates and kin are displayed by a fascinating, but limited, subset of species across the animal kingdom (Kleiman, 1977; Pfennig and Sherman, 1995; Robinson et al., 2019). Such attachments, which form the basis of diverse and complex social systems, are observed in species that have evolved the capacity to form lasting bonds between individuals, suggesting that they are innate with a strong underlying genetic component (Insel and Young, 2000; Maruska, 2015; Staes et al., 2021; Wilson and Hölldobler, 2005). Progress in understanding the molecular or neural networks that promote social attachment has been hindered because traditional genetic model organisms such as *C. elegans, D. melanogaster, D. rerio*, and *M. musculus* do not display enduring attachments as adults. Here we report the use of a CRISPR-based mutagenesis in prairie voles *(Microtus ochrogaster)* to probe signaling pathways previously implicated in social attachment.

Adult prairie voles exhibit social attachment behavior such that mating partners form an enduring bond with each other. This attachment behavior, commonly referred to as pair bonding, has been observed in ethological studies in the wild as well as in the laboratory setting (Carter and Keverne, 2002; Getz et al., 1981). Pair bonded voles spend time together in close proximity (huddling behavior) and show a social preference for each other over a potential new mating partner. Neuropeptide signaling has long been known to control the display of social behaviors across diverse species (Donaldson and Young, 2008; Garrison et al., 2012; Insel et al., 1995a, 1998; Wagenaar et al., 2010). Differences in neural populations regulated by these pathways correlate with interspecies variations in social structure (Ophir et al., 2012; Shapiro and Insel, 1992; Winslow et al., 1993). Intriguingly, the evolution of varied complex social systems and affiliative behaviors, including social monogamy, has repeatedly converged upon the nonapeptide hormones oxytocin (Oxt) and arginine vasopressin (Avp) and their orthologs across phylogenies (Klatt and Goodson, 2013; Nowicki et al., 2020; O’Connor et al., 2016).

Ethological studies, using live-trapping of wild prairie voles, reported that mating pairs are more likely to be trapped together than is expected by chance (Getz et al., 1981). Similar studies of closely related species such as meadow voles (*M. pennsylvanicus*), that do not pair bond, showed that live-traps contained single animals (Getz et al., 1981). Subsequent studies demonstrated that pair bonding behavior can also be observed in a laboratory setting (Cho et al., 1999; Insel and Hulihan, 1995). Pair bonding is displayed as a suite of behavioral traits, the most commonly measured of which is a preference for the familiar partner over a novel stranger. Pair bonded animals prefer to huddle with their partners compared to exploring unfamiliar conspecifics of the opposite sex (Williams et al., 1992). Such partner preference is also accompanied by aggression toward such unfamiliar conspecifics of the opposite sex, indicative of active rejection of potential new mates (Wang et al., 1997).

Comparative studies between socially monogamous and non-monogamous vole species revealed striking differences in OxtR expression in brain regions thought to be important for social attachment, and implicated natural variation within species in specific aspects of pair bonding and attachment behaviors (Heinrichs et al., 2009; Insel et al., 1994; King et al., 2016; Ophir et al., 2012; Shapiro and Insel, 1992; Walum and Young, 2018). Pharmacological studies from multiple groups have shown that oxytocin (Oxt) is sufficient to induce pair bonding behavior in otherwise naive voles and the administration of OxtR-antagonists causes the loss of these behaviors (Carter et al., 2008; Cho et al., 1999; Insel et al., 1995a; Winslow et al., 1993). Viral manipulations of OxtR expression in specific brain regions of prairie voles also recapitulate findings from such pharmacological studies (Keebaugh and Young, 2011; Keebaugh et al., 2015). Taken together, these findings suggest a critical role for Oxt signaling via its cognate receptor OxtR in driving pair bonding behaviors in this species.

Prairie voles, like many other animals that display pair bonding behavior, exhibit biparental care of their young, and oxytocin signaling is thought to control these behaviors as well (Keebaugh et al., 2015; Ophir et al., 2013). Oxytocin is additionally critical for milk let-down, the reflexive release of milk triggered by sensory stimuli associated with suckling (Lee et al., 2008; Young III et al., 1996). All pups born to female mice null for Oxt or OxtR die shortly after birth because of complete failure of milk let-down (Nishimori et al., 1996; Takayanagi Yuki et al., 2005). In addition to this role in lactation, stimulation of Oxt-expressing neurons in virgin female mice induces pup retrieval behaviors typical of lactating females (Marlin et al., 2015). Hence, decades of research implicate both the neuropeptide and its receptor in a large repertoire of parenting behaviors.

To test the genetic requirement of OxtR in pair bonding and parental behaviors, we employed a CRISPR-based approach to generate mutant prairie voles null for this receptor. Surprisingly, male and female prairie voles homozygous for each of the three distinct loss-of-function OxtR alleles displayed pair bonding. Moreover, we observed that OxtR null females were capable of raising pups to weaning. We therefore conclude that, contrary to previous assumptions, OxtR function is dispensable for parental behaviors and pair bonding.

## Results

### CRISPR mutagenesis can reliably generate multiple null alleles of *OxtR* in prairie voles

To perform CRISPR-based gene targeting in prairie voles, we developed a protocol to obtain single cell embryos for injection of Cas9 ribonucleoprotein complexes. We were unable to achieve successful superovulation using hormonal supplementation protocols including those previously reported (Horie et al., 2019), and therefore we implemented a timed-mating strategy to harvest embryos synchronized at specific developmental time point. We harvested 0.5 day single cell embryos using this protocol, an approach that yielded 3.7±1.2 embryos/female.

Because we maintain prairie voles in an outbred background, we were concerned about natural variation in OxtR coding sequence that could potentially reduce targeting efficiency by specific small guide RNAs (sgRNAs). Indeed, we observed 3 synonymous substitutions in exon 1 of the *OxtR* locus from 4 voles in our colony (Figure S1A). We designed 8 protospacer adjacent motif (PAM) site-anchored sgRNAs based on conserved sequences in exon 1. We initially tested whether these sgRNAs could generate mutations in exon 1 *in vitro*, injecting them as Cas9 ribonucleoprotein complexes individually into single cell embryos and genotyping 4 days later at the blastocyst stage (Figures S1B-D). Two sgRNAs consistently yielded mutations in exon 1, and sequencing revealed multiple mutations in the same blastocyst, suggesting that CRISPR-targeting had occurred after the single cell stage (Figures S1E-F).

**Figure 1.**
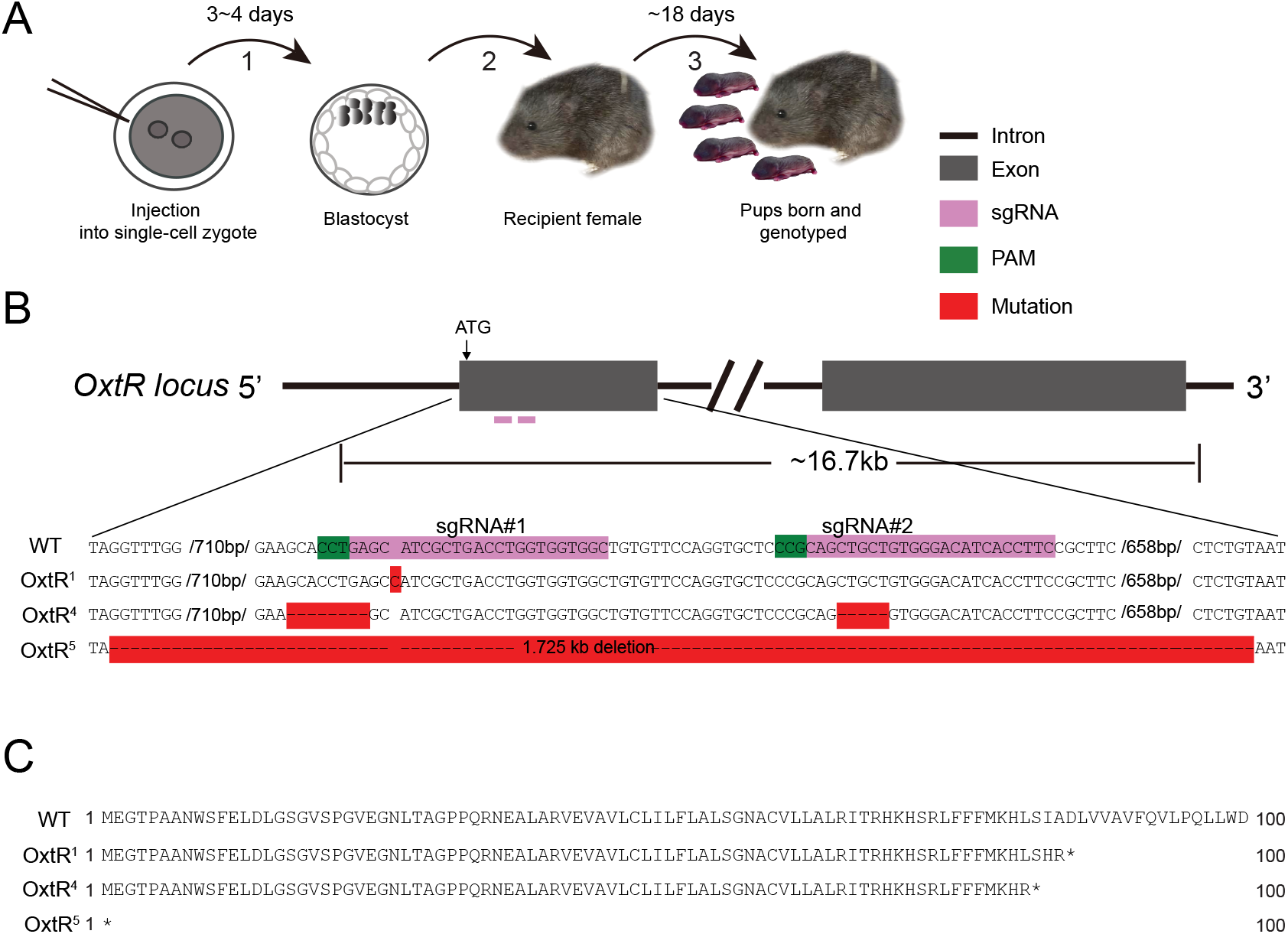
CRISPR mutagenesis can reliably generate multiple null alleles of *OxtR* in prairie voles. **A.** Schematic of CRISPR-based targeting to generate OxtR mutant prairie voles. Single cell embryos injected with Cas9-sgRNA ribonucleoprotein were cultured (1) to the blastocyst stage and transferred (2) to pseudopregnant recipient females who carried the embryos to term (3). **B.** Schematic (top) of Oxtr locus encompassing first two exons. DNA sequence of WT and targeted *OxtR* alleles. Dash/missing nucleotide represents a deletion (Oxtr^4^, Oxtr^5^) and red highlighted “C” (Oxtr^1^) is an insertion. PAM, protospacer adjacent motif. **C.** Predicted amino acid sequence of WT OxtR, OxtR^1^, OxtR^4^, and OxtR^5^ (only first 100 amino acids shown).

We proceeded to co-inject these two sgRNAs into single cell embryos, cultured them *in vitro* to the blastocyst stage, and then transferred them into recipient pseudopregnant prairie voles (Figure 1A). Each recipient female gave birth to 1-2 pups (~10% of embryos transferred) ~16 days following embryo transfer. Given the significant chimerism we observed following genotyping of blastocysts injected with our chosen sgRNAs (Figures S1E-F) and the low yield of liveborn pups following embryo transfer, we mated each founder (G0) to a wildtype (WT) partner even when genotyping tail samples from these founders revealed no mutations. We generated founders carrying 3 distinct *OxtR* alleles that transmitted via the germline (Figures 1B,C). *OxtR^1^* has a 1 base pair (bp) insertion that is predicted to yield an 84 amino acid (aa) peptide, *OxtR^4^* contains 2 small deletions that are predicted to yield an 81 aa peptide, and *OxtR^5^* contains a large deletion extending 1.5 kbp beyond the sequence targeted by the two sgRNAs. To minimize carry-over of potential off-target mutations in our lines, we independently outcrossed all lines of each of the three mutant alleles to WT voles in our colonies.

### CRISPR-generated mutations in OxtR produce loss of function alleles

Each of the three *OxtR* alleles we generated is predicted to generate non-functional proteins (Figure1C). We tested this directly by performing a ligand binding assay *in situ*, using a radiolabeled small molecule competitive agonist for OxtR. These studies showed a complete lack of binding in homozygous null mutants of both sexes of *OxtR^1^* (Figure 2), *OxtR^4^*, and *OxtR^5^* (Figure S2), demonstrating absence of all ligand-binding OxtR *in vivo*. Consistent with the lack of requirement for OxtR for survival, we observed Mendelian ratios of WT, heterozygous, and homozygous pups born to heterozygous parents (OxtR^1^ 29:60:24, OxtR^4^ 37:62:32 and OxtR^5^ 25:55:26).

**Figure 2.**
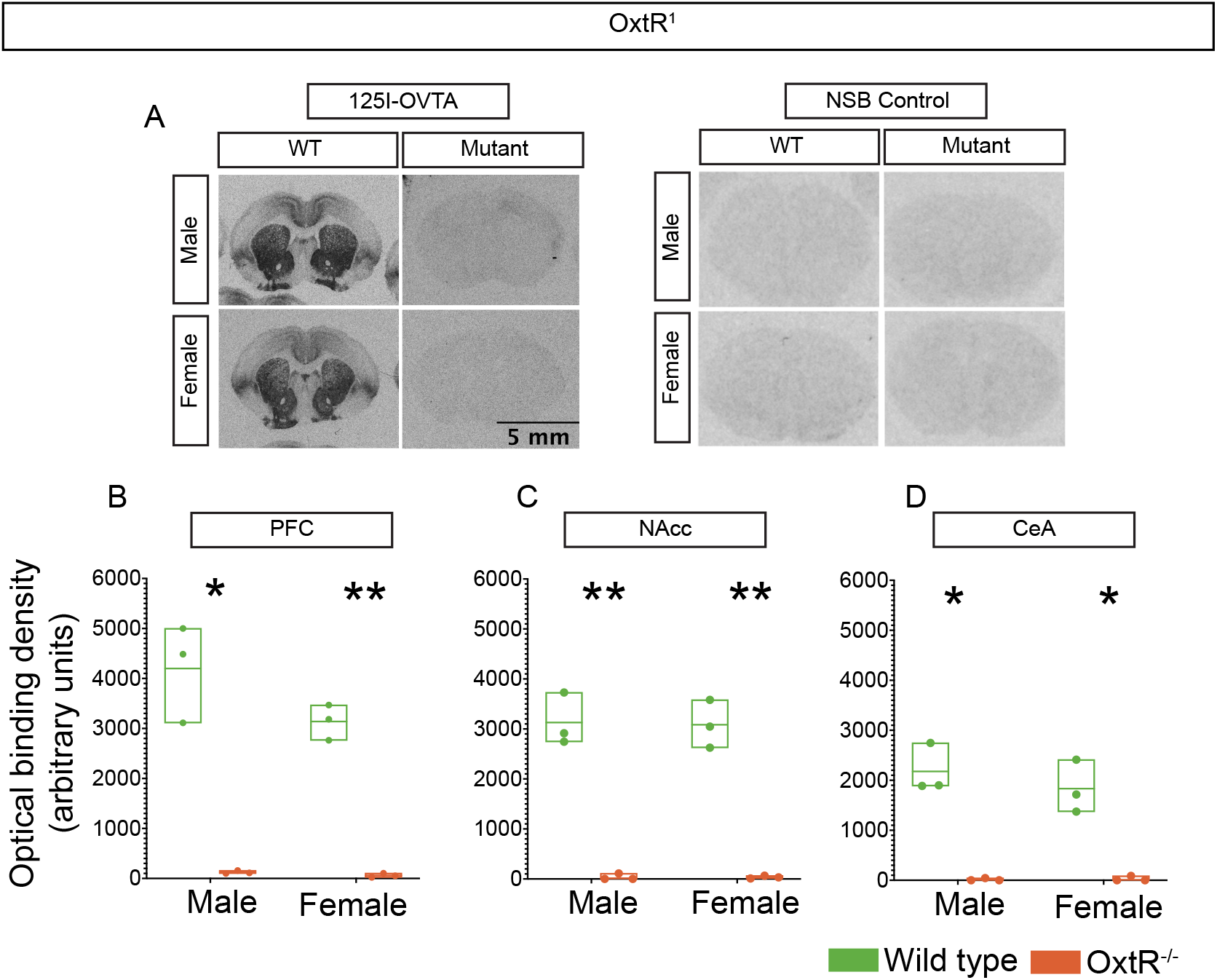
CRISPR-generated mutations in OxtR produce loss of function alleles. **A.** Loss of binding with the competitive agonist 125I-OVTA visualized in coronal sections through the rostral telencephalon of OxtR^1^ homozygous mutant voles (A). **B-D.** Optical density-based quantification of binding to 125I-OVTA shows that binding is essentially undetectable in mutants null for OxtR^1^ in PFC (prefrontal cortex) (B), NAcc (nucleus accumbens) (C), and CeA (central amygdala) (D). Scale bar =5 mm; Box plot Min to Max, mid-line denotes mean; n = 3 (B, C, D) *p<0.05, **p<0.01, ***p<0.001.

### Female and male prairie voles lacking OxtR exhibit robust pair bonding behaviors

Prairie voles are induced ovulators, and following a short period of cohabitation between opposite sex animals, co-housed pairs will mate and subsequently display pair bonding (Insel et al., 1995b; Williams et al., 1992). To test whether our null mutants display deficits in pair bonding, we co-housed sexually naive males and females for 7 days, a period previously shown to be sufficient to induce pair bonding (Insel et al., 1995b; Williams et al., 1992). WT or homozygous mutants were paired with an unfamiliar, unrelated WT animal of the opposite sex. We used two different paradigms to assay partner preference and behaviors between the experimental subject and a cohoused partner or novel unfamiliar conspecific of the opposite sex. In the 3 chamber assay, the experimental animal, its pair bonded partner, and an unfamiliar, opposite-sex, conspecific are housed in separate interconnected chambers such that only the experimental subject has free access to all chambers (Figure 3A) (Cho et al., 1999). Alternatively, we used a single, partially-divided chamber in which the cohoused partner and unfamiliar conspecific are tethered at opposite ends from each other, with the experimental subject free to exhibit affiliative or aggressive behaviors toward either tethered animal (Figure 3B) (Beery and Shambaugh, 2021; Beery et al., 2018).

We tested pair bonding of OxtR^1^ mutants and their control group in the partially divided chamber paradigm and that of OxtR^4^ and OxtR^5^ in the 3-chamber assay. Surprisingly, we observed that males and females homozygous for each of the three, null, mutant *OxtR* alleles all spent more time huddling with their partners and displayed aggression towards the unfamiliar conspecific of the opposite sex, in a manner comparable to WT animals regardless of the apparatus or setting used to test these behaviors (Figures 3C-H, S3). Thus, we find that prairie voles demonstrate pair bonding behaviors in the absence of OxtR.

**Figure 3.**
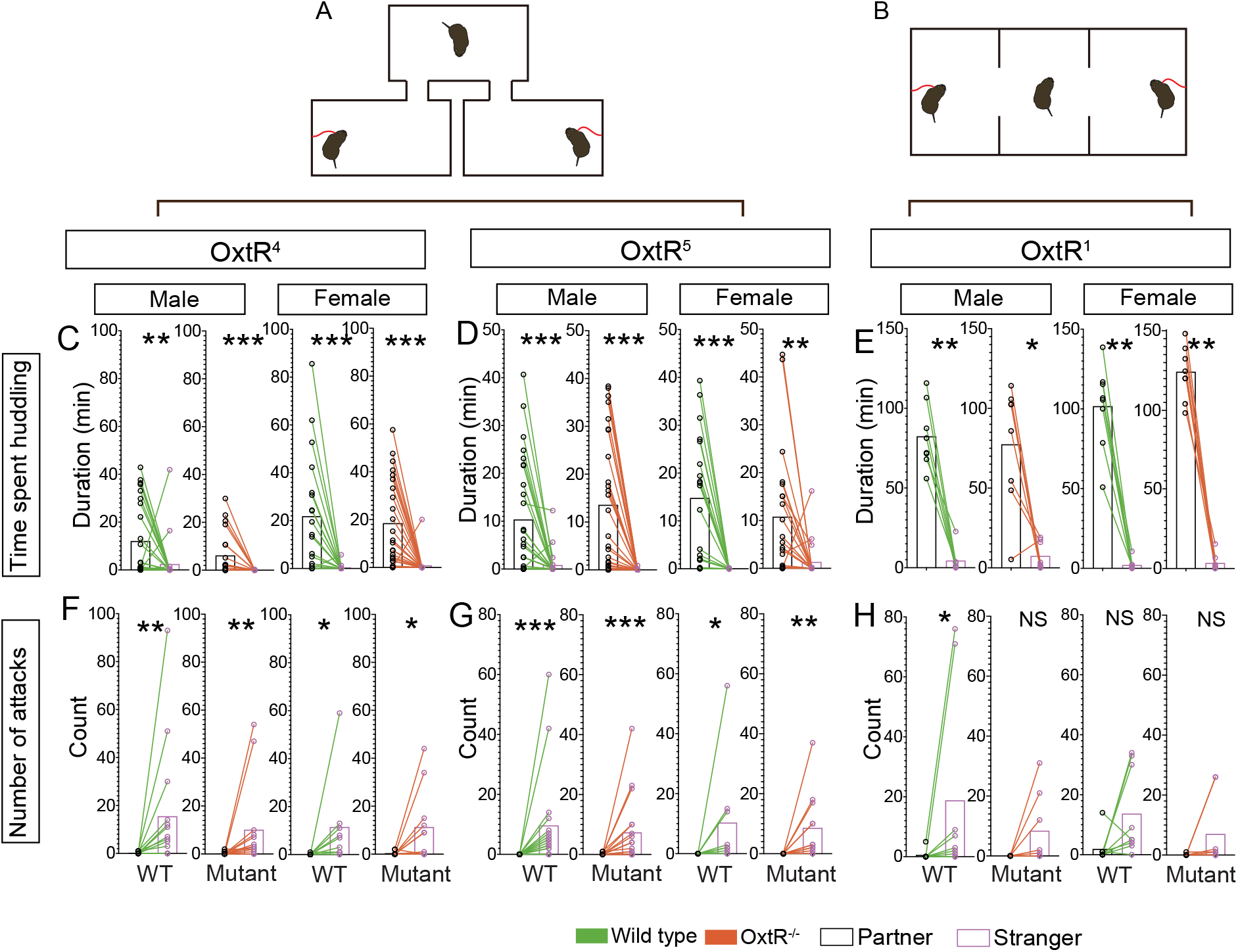
Female and male prairie voles lacking OxtR exhibit robust pair bonding behaviors. **A-B.** Schematic of partner preference test performed in a 3-chamber (A) or single chamber with two partitions (B). Experimental vole is free to move between chambers (A) or partitioned space (B) whereas stimulus voles (opposite sex partner and stranger) are restrained by tethers (red). **C-E.** WT, OxtR^4-/-^ homozygous mutant (C), OxtR^5-/-^ homozygous mutant (D), and OxtR^1-/-^ homozygous mutant (E) voles spent more time with their partner than a stranger of the opposite sex. **F-H.** WT, OxtR^4-/-^ (F), OxtR^5-/-^ (G) and OxtR^1-/-^ (H) null mutants attacked the stranger more frequently than their partner. Mean ± SEM; n = 25 WT and 21 mutant males, 19 WT and 21 mutant females (C); 27 WT and mutant males each, 19 WT and 21 mutant females (D); 8 WT and 8 mutant males, 8 WT and 8 mutant females each (E); 12 WT and mutant males each, 8 WT and 11 mutant females (F); 15 WT and mutant males each, 8 WT and 11 mutant females 15 (G); 8 WT and mutant males and females each (H); *p<0.05, **p<0.01, ***p<0.001; N.S., not significant.

### Prairie voles lacking OxtR bear viable pups and display bi-parental care similar to wild types

Given the concordance in pair bonding displays by animals homozygous for each of the three mutant *OxtR* alleles, we tested a subset of these for performance in parenting. *OxtR^4^* or *OxtR^5^* homozygous mutants and their WT siblings were paired with WT animals of the opposite sex until parturition and then tested for parental care of their progeny. Both male and female prairie vole parents interact intensively with their pups, spending prolonged periods huddling, licking, and grooming them. Experimental interference during the pre-weaning period can disrupt both subsequent displays of parental care as well as the repertoire of adult social behaviors exhibited by the pups (Bales et al., 2007a; Danoff et al., 2021). Accordingly, the standard assay is to observe the duration of parental interactions with their pups in the first few days after parturition and document pup survival at weaning (Perkeybile et al., 2013).

We observed that OxtR null parents interacted equivalently with their pups compared to their WT counterparts (Figures 4A-F). Both WT and mutant parents spent the majority of their time in the nest, in direct contact with their litters, and, in the case of mothers, nursing pups. Unlike mice lacking OxtR (Takayanagi Yuki et al., 2005), we never observed cages in which pups were scattered across the cage floor, indicative of effective retrieval to the nesting area of pups who had wandered away.

**Figure 4.**
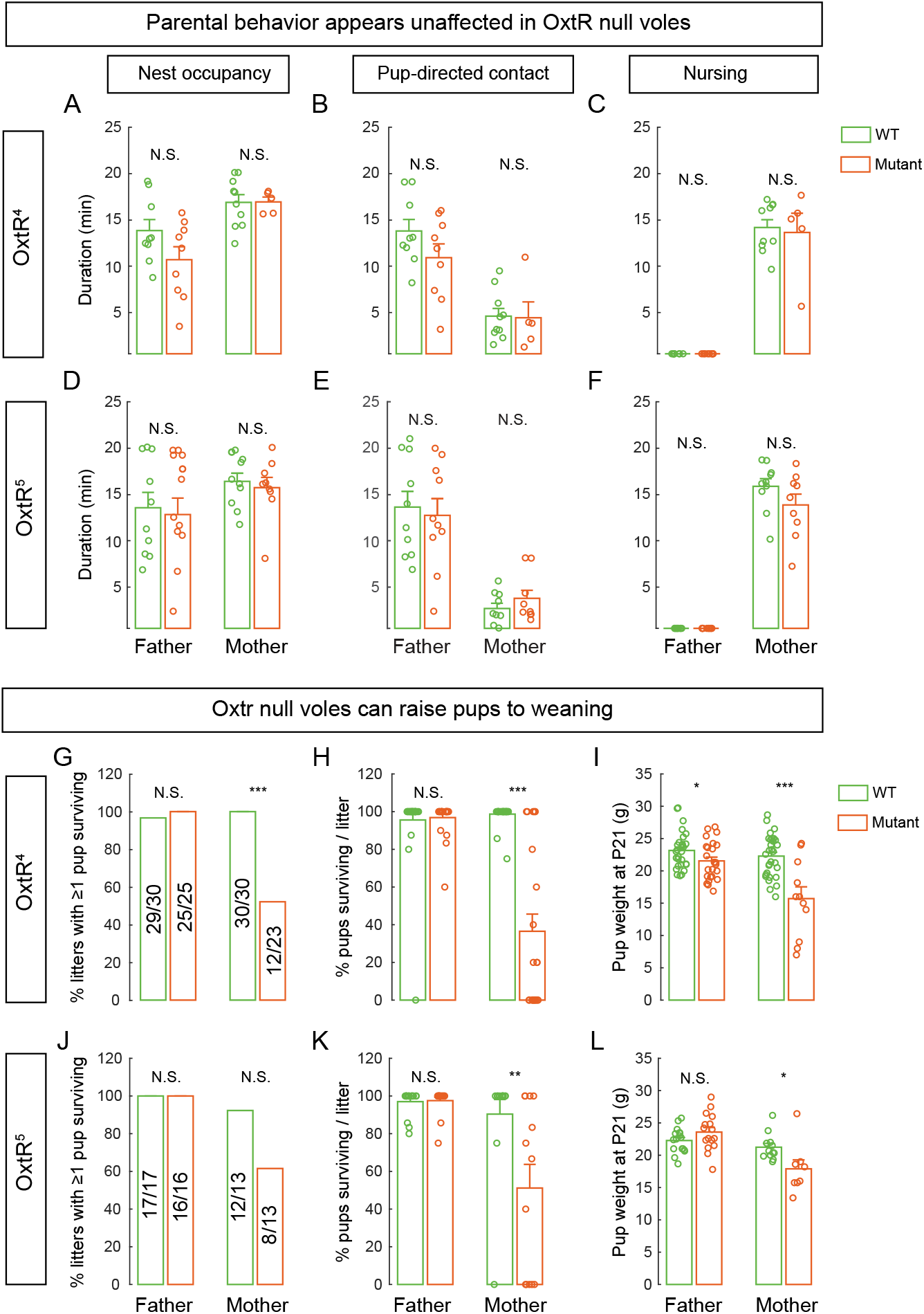
Prairie voles lacking OxtR bear viable pups and display bi-parental care similar to wild types. **A-F.** WT, OxtR^4-/-^, and OxtR^5-/-^ mothers and fathers exhibit equivalent nest occupancy (A, D), pup-directed contacts (B, E), or nursing behavior (C, F) with their pups. **G, J.** OxtR^4^ mothers wean fewer litters compared to WT mothers. **H, K.** Litters from OxtR^4-/-^ and OxtR^5-/-^ mothers have fewer pups surviving to weaning compared to WT mothers. Only litters with ≥1 pup surviving to weaning were analyzed. **I, L.** OxtR^4-/-^ mothers or fathers and OxtR^5-/-^ mothers weaned pups with significantly lower body weight compared to WT parents. Mean ± SEM; n = 9 WT and mutant males each, 10 WT and 5 mutant females (A-C); 10 WT and mutant males each, 10 WT and 9 mutant females (D,F); 10 WT and mutant males each, 9 WT and mutant females each (E); *p<0.05, **p<0.01, ***p<0.001; N.S., not significant.

Surprisingly, we noticed that pups born of mothers homozygous for either of two OxtR alleles survived past the first few hours of birth and many weaned successfully, suggesting that mutant females could nurse some of their young (Figures 4G,H,J,K). *OxtR^4^* null mothers had significantly fewer surviving litters at weaning compared to WT mothers (Figure 4G). *OxtR^5^* null mothers also had fewer surviving litters at weaning, although this did not reach statistical significance (Figure 4J). Of the litters that survived to weaning to experimental females, *OxtR* null mothers had fewer pups per litter (Figures 4H,K). Pups born to mutant mothers at weaning weighed significantly less than pups born to WT mothers, suggesting a defect in milk let-down or subtle deficits in nursing behavior (Figures 4I,L). Consistent with a deficit in milk let-down, gross examination of the nipple area revealed significantly less engorgement in females mutant for OxtR compared to WT mothers (not shown).

Both WT and mutant fathers successfully raised their litters to weaning (Figures 4G,H,J,K). Pups raised by *OxtR^4^*, but not *OxtR^5^*, null fathers weighed less than those raised by WT fathers, suggesting a role for OxtR in fathers in their ability to raise thriving pups. Our findings show that, in prairie voles, *OxtR* function is dispensable for biparental care and can be partially bypassed for milk let-down.

## Discussion

CRISPR-mutagenesis has been successfully used to manipulate a large variety of species of animals, allowing scientists the freedom to establish new model organisms for biological phenomena that are not observed in worms, fruit flies, zebrafish, or mice. We have utilized CRISPR-based targeting to test the role of the OxtR locus in prairie voles, given the phylogenetically conserved role of this pathway in modulating attachment and social behaviors (Feldman et al., 2016). We find that pair bonding behaviors, nursing, and successful weaning of pups can all occur in the complete absence of OxtR-mediated signaling. Our findings, consistent across multiple paradigms, three labs, and three null alleles of *OxtR*, are in contrast to prior studies that highlight the importance of OxtR signaling in pair bonding and parental behaviors in prairie voles (Bales et al., 2004; Cho et al., 1999; Insel et al., 1995b, 1998; Keebaugh and Young, 2011; Keebaugh et al., 2015; King et al., 2016). A key difference between our work and preceding studies is that the latter were conducted in adult animals, using pharmacological or viral misexpression strategies to determine a role for OxtR function in behavior. The studies we present illustrate the importance of complementary pharmacological and genetic approaches to understand the functional relevance of signaling pathways *in vivo*.

Despite efforts to demonstrate the specificity of pharmacologic approaches to manipulate OxtR activity, it remains possible that the compounds employed in these studies alter yet-to-be-identified pathways that contribute to pair bonding. In such a model, exogenous ligands that activate OxtR bind to and stimulate additional receptors, while compounds that antagonize Oxt binding to OxtR could also inhibit either its binding to additional receptors or other ligand-receptor interactions (Bales et al., 2007b; Chini and Manning, 2007). It is possible that compensatory pathways activated due to constitutive loss of function of OxtR in our studies obscure a functional role for this receptor in pair bonding and parenting (El-Brolosy and Stainier, 2017; Teng et al., 2013). We did not find an increase in Oxt or Avp expression (Figure S4) indicative of such compensation, though we cannot exclude more subtle changes in the expression of these neuropeptides or other pathways that substitute for the loss of OxtR signaling.

Future studies with targeted loss of OxtR in specific brain regions of adult voles will reveal whether it is essential for these behaviors. Alternatively, OxtR and Avpr1a signaling might redundantly regulate pair bonding and parenting; Oxt can also signal via Avpr1a and it may therefore modulate these behaviors entirely through Avpr1a rather than OxtrR(Anacker et al., 2016; Chini and Fanelli, 2000). These possibilities can be disambiguated by testing the behavioral performance of voles null for Avpr1a or Avpr1a and OxtR. Determination of the non-OxtR targets of the pharmacological agents that have successfully abolished pair bonding behaviors might also lead to insights into the underlying neural circuits and molecular signaling pathways. In any event, our studies rigorously demonstrate that, in three different loss of function mutations generated and tested independently, OxtR signaling is not genetically required in male and female prairie voles for either pair bonding or multiple aspects of parenting.

Female mice null for OxtR show a complete failure of milk let-down and nursing behavior such that none of their pups survives beyond the day of birth (Takayanagi Yuki et al., 2005; Wakerley et al., 1973; Young III et al., 1996). In contrast, many pups born to prairie vole mothers lacking OxtR survive to weaning, albeit with reduced weight. It is possible that this phenotypic difference between mice and voles mutant for OxtR is a consequence of rapid evolution of pathways that regulate nursing and parenting in mammals (Feldman et al., 2016; Paré et al., 2016; Staes et al., 2021). Alternatively, this difference in phenotypes could reflect the fact that the OxtR mutant mice were maintained on an inbred background whereas our colony is consistently outbred to wild voles (Crawley et al., 1997; Simpson et al., 1997). It will be interesting to determine whether OxtR null mice on an outbred, wild background can also exhibit parental behaviors. If true, this would suggest that inbreeding has led to reliance on OxtR signaling for parenting in mice rather than a fundamental difference in species-specific roles for OxtR in parental displays.

Oxt signaling has also been implicated in affiliative displays in humans suggesting a conserved role for this neuropeptide hormone in these social behaviors across ~90 million years of evolution (Walum and Young, 2018). Based on observations in prairie voles and other mammals, including humans, clinical trials have used exogenous oxytocin or small molecule ligands to OxtR to ameliorate the deficits in social attachment and cognition seen in multiple psychiatric conditions; these studies however have yielded mixed results (Di Simplicio and Harmer, 2016; Feifel et al., 2010; Green and Hollander, 2010; Heinrichs et al., 2009; Rubin et al., 2010; Sikich et al., 2021). Regardless of compensatory mechanisms that might mitigate the loss of OxtR-signaling, our observation that OxtR signaling is not required genetically for pair bond formation or parenting in prairie voles suggests that we require a more refined understanding of the molecular pathways underlying social attachment behaviors. New genetic models like prairie voles may be more fitting for the pursuit of testing therapeutic interventions for the diverse array of conditions that affect attachment and social behavior, as such whole-animal mutants better represent what may occur in patients with mutations associated with neuropsychiatric disorders.

## Acknowledgments

The authors would like to thank members of the Bales lab for assistance and advice in animal husbandry and the Bales, Shah, and Manoli labs for helpful discussion and comments on the manuscript. In addition, we would like to thank Jake Freimer, Ron Parchem, Archana Shenoy, and the Blelloch lab for help and advice regarding embryo harvesting and manipulation, and the Doudna, Jaenisch, and Zhang labs for advice regarding CRISPR mutagenesis.

## Author Contributions

Conceptualization, K.L.B., D.S.M. and N.M.S.; Methodology, A.B., K.L.B., K.M.B., M.A.M., D.S.M., F.D.R., N.M.S., R.S., Y.C.W.; Investigation, J.M.B., K.M.B., I.E., J.K., R.L., M.A.M., S.P., F.D.R., T.C.S., A.H.M.S., R.S., Y.C.W.; Visualization, K.M.B., M.A.M., R.S., Y.C.W.; Funding acquisition, K.L.B., D.S.M., N.M.S.; Writing –Original Draft, K.M.B., R.S.; Writing – review & editing, K.L.B., K.M.B., D.S.M., M.A.M., N.M.S., R.S., Y.C.W.

## Star Methods

### Experimental model and subject details

#### Animals

Subjects were laboratory-bred prairie voles (*Microtus ochrogaster*) which originated through systematic outbreeding of a wild stock captured near Champaign, Illinois. Prairie voles were bred, maintained, and tested at three distinct sites: University of California San Francisco, University of California Davis, and Stanford University. Sexually naïve male and female animals were group weaned at 21 ± 1 days and separated to pair-housing with either a same-sex sibling or an age-matched same-sex non-sibling (about half to each type of pairing). Voles were maintained under a 14:10 h light-dark cycle in clear plastic cages (45 × 25 × 15 cm) with bedding, nesting material (nestlet), and a PVC hiding tube. Rooms were maintained at approximately 20°C, and food and water were available *ad libitum*.

Breeding pairs were established between two heterozygotes or a homozygous OxtR mutant male and heterozygous female partner from a breeding line maintained by the respective labs. Pup weights were recorded at birth and at weaning date. Voles were randomly assigned into experimental groups when they reached 7-9 weeks of age at the start of testing. This study was carried out in accordance with the recommendations in the Guide for the Care and Use of Laboratory Animals published by the National Research Council. The protocol was approved by the Institutional Animal Care and Use Committee at each respective institution.

### Method details

#### Isolation of prairie vole embryos for gene targeting

Unlike mice, ovulation in prairie voles is behaviorally induced. All attempts to artificially super-ovulate prairie voles yielded poor results in our hands, differing from findings of a prior study (Horie et al., 2019). We therefore developed a timed mating protocol where we placed 6-8 week old females with age matched males for 12 hours and then placed a physical barrier between them and housed them for 3 another 24 hours. We empirically determined that this induced estrus in all the females we paired, and the females all yield fertilized embryos approximately 18 hours after the removal of the barrier.

#### Mutagenesis and embryo manipulation for gene targeting

We obtained the genomic sequence of exon 1 of OxtR from 4 randomly chosen animals from our colony and identified polymorphisms that correspond to 3 synonymous substitutions. Based on these sequences, we designed 8 sgRNAs targeting exon 1, and adopted microinjection protocols commonly used to manipulate embryos from mice to test varying concentrations of the guides along with either 20 ng/ul or 100ng/ul of Cas9 mRNA based on studies in mice, rats, and other rodent species (Yang et al., 2013). We cultured embryos to the blastocyst stage, with ~45% of embryos surviving following injection. We then harvested genomic DNA from individual blastocysts, and assayed for mutagenesis of OxtR by Surveyor PCR, and sequencing of genomic PCR products (Ran et al., 2013). These analyses revealed significant mosaicism in blastocysts, suggesting that mutagenesis in vole embryos occurred significantly later than the first division. Based on these studies, we identified 2 sgRNAs (sgRNA1: GAGCATCGCTGACCTGGTGGTGGC, sgRNA2: CAGCTGCTGTGGGACATCACCTTC) that reliably yielded detectable mutations in embryos and selected these for further use. We performed pronuclear injections of the 2 sgRNA guides, 2μl Cas9 protein (PNA BIO, INC CP02, 5μg/μl) and 2μl cas9 mRNA (TRILINK BIOTECHNOLOGIES, INC L-7206, 1μg/μl) and cultured the manipulated embryos to the blastocyst stage. We generated pseudopregnant recipient females using our timed mating protocol with vasectomized males and surgically implanted the manipulated embryos into their oviduct. Using such an approach, we developed a protocol whereby ~10 embryos are transferred to each uterine horn to obtain 3-5 live born pups per female.

#### Isolation and Outcrossing of OxtR Alleles

Tail samples were taken from all of the founders (G0s) and their progeny (F1s) following pairing with wild type mates. If these tail samples yielded a mutation in the *OxtR* locus, the animals were then paired with wild types of the opposite sex and their pups (F2s) were genotyped by PCR of the OxtR locus to verify persistent germline transmission of the mutant allele. For subsequent outcrosses, at each generation (Oxtr^4^ and Oxtr^5^, 3 generations; OxtR^1^, 7 generations), following PCR genotyping for the OxtR allele, a heterozygous male and female of each allele were independently paired with an WT of the opposite sex to isolate OxtR alleles from any off-target effects of CRISPR mutagenesis.

#### Autoradiography

Wildtype and homozygous mutant siblings were sacrificed using CO2 and their brains were dissected and rapidly frozen on powdered dry ice before storing at −80°C. 20μm sections were thaw mounted onto super mount frost plus slides. Slide mounted sections were thawed until dry and fixed for two minutes in 0.1% paraformaldehyde (0.1% paraformaldehyde in 0.1M PBS). Slides were rinsed 2×10 minutes in 50mM Tris pH 7.4, then incubated for 90 minutes at room temperature in a solution (50 mM Tris, 10 mM MgCl2, 0.1% BSA, 0.05% bacitracin, 50 pM radioligand) containing the radioactively labeled 125I-ornithine vasotocin analog vasotocin, d(CH2)5[Tyr(Me)2,Thr4,Orn8,(125I)Tyr9-NH2] (125I-OVTA, PerkinElmer, Inc.). Slides processed for non-specific binding were incubated with an additional 50 uM non-radioactive ligand [Thr4Gly7]-oxytocin (Bachem). All slides were washed 3×5 min in chilled Tris-MgCl2 (50mM Tris, 10mM MgCl2, pH 7.4), followed by a 30-minute soak in Tris-MgCl2 on ice. Slides were quickly dipped in cold, distilled water and then air dried and exposed to Kodak BioMax MR film (Kodak, Rochester, NE, USA) for 3 days and subsequently developed. To quantify ligand binding, we first defined anatomical centers using the Allen Mouse Brain Atlas.

The 125I-OVTA binding was quantified directly (measured as optical binding density, OBD) from the film using a light box, a top mounted camera, and the MCID Core Digital Densitometry system (Cambridge, UK) according to methods previously described (Rogers et al., 2021). Values for OBD from a set of 125l autoradiography standards (American Radiolabeled Chemicals, Inc., St. Louis, MO, USA) were loaded into MCID in order to generate a standard curve, from which 5 OBD values for each ROI were extrapolated (Oxtr^4^ and Oxtr^5^). Average OBD values were calculated for each ROI within each individual specimen, as well as for one background area where no binding was detected, which served as a measure of non-specific binding. This average background/non-specific binding was subtracted from the ROI measurements to yield normalized OBDs across specimens and correct for individual variation in non-specific binding across individuals. Autoradiography performed on Oxtr^1^ was calculated with respect to the background before adjusting the background to zero. Non-standard binding controls performed on Oxtr^1^ showed that there was no difference between signal observed in the mutant and background. The signal intensities were averaged across males and females of both wild type and mutant siblings from a particular brain region and compared using the student’s T test.

#### Partner preference assay

The partner preference test (Williams et al., 1992) was used to assess pair bond formation (Ahern and Young, 2009). Subjects were cohoused with an opposite-sex, wildtype animal (partner). Analysis of partner preference between wild type siblings from the OxtR^1^ background and wild type animals from this colony showed no difference on the partner preference test. Pairing of OxtR^1^ was followed by a timed mating protocol for synchronization of estrous in which paired animals are separated by a clear, plastic barrier 18 hours following pairing. The barrier was left in place for 24 hours and then removed allowing the subject free access to the partner animal thereafter. Regardless of which laboratory performed testing, all animals were allowed to cohabitate for one week prior to behavioral testing. Two different, well-established apparatuses were used to test partner preference. In the three-cage design used to test Oxtr^4^ and Oxtr^5^, the subject, their partner, and an unfamiliar opposite-sex conspecific (stranger) were placed in a three-chamber apparatus made of three small polycarbonate cages (27×16×16 cm) joined by clear Plexiglas tubes. Strangers were selected to be of similar age and size as the partner animal. Both the partner and the stranger were tethered in two separate, end cages, and the test subject was placed, untethered in a central cage. The test subject was free to access any of the three cages over a period of 3 hours. All behaviors were video recorded from an approach centered on the two end cages. In the apparatus design used to assay OxtR^1^ animals, the subject, partner, and an opposite sex stranger are placed in a three-chamber apparatus with open top and 10 x 32 in walls. The partner and stranger animals are tethered on either end of the three-chamber arena and the subject allowed free access, again over a period of 3 hours. Behaviors are recorded from a top view camera capturing the entire apparatus. Videos were scored post-test by validated scorers blind to condition. Observed behaviors included location (i.e., duration of time in partner, stranger, and neutral cages), duration of stationary huddling or >50% side-by-side contact between the partner and stranger animals, and frequency of aggressive behavior (i.e., lunges).

#### Parental behavior

A total of 80 individuals, distributed across sex (male or female), strain (Oxtr^4^ or Oxtr^5^) and genotype (wild-type or mutant) were paired with a wild type colony animal of the other sex. Following partner preference testing, pairs were left in large polycarbonate cages (44×22×16cm) with sterilized cotton for nesting material. All litters from 21 breeder pairs were observed on two separate days in the morning (08:00-11:00) and two separate days in the evening (15:00-18:00) for a total of four, 20-minute focal observations during the neonatal period (P1-P3). Parenting behavior was observed according to methods previously outlined (Perkeybile et al., 2013). Following the birth of pups, breeding pairs were observed for a suite of pup-directed behaviors on two separate days in the morning and in the afternoon for a total of four, 20-minutes focal samples between post 7 natal days 1 and 3. Eight individuals were not observed because of their death or the death of their partner (2 sires, 6 dams). Notably, of the two sires, one was wild type and one was mutant; and of the six dams, all six were mutant, split between OxtR^4^ (n = 5) and OxtR^5^ (n = 1). Increased rates of culling or natural death may have been due to reproductive complications in which oxytocin plays a significant role (pregnancy, birthing, and lactation). All observations were made in the home cage. Each parent was observed and left undisturbed during each 20-minute observation. The duration (seconds) of each behavior was recorded, and durations of all direct parenting behaviors (e.g. licking/grooming, huddling, etc.) were recorded within each observation period. All behaviors were live recorded using behavioral software (www.behaviortracker.com,) or ScoreVideo (matlab).

#### RT-quantitative PCR

Adult WT and OxtR mutant voles were deeply anesthetized by IP injection with 2.5% Avertin and euthanized by decapitation. The brain was quickly dissected out and sectioned into 500 μm coronal slices using a brain matrix mold (BrainTree Scientific) chilled on ice. Slices were floated in ice-cold phosphate buffered saline (PBS) and the paraventricular nucleus (PVN) was identified using anatomical landmarks and dissected using a Zeiss microscope. For Oxtr^4^, tissue was dissected and RNA purified from individual animals. For OxtR^5^, tissue from 2-3 animals was pooled prior to RNA extraction. Tissue was flash-frozen on dry ice and stored at −80°C until further processing. Total RNA was extracted using TRIzol according to manufacturer’s instructions and quantified using NanoDrop (Thermo Scientific). We used 5 μl total RNA to program a 20 μl reverse transcription reaction using the ProtoScript cDNA synthesis kit with random hexamer priming. We used 2μl of the RT reaction to run real-time quantitative PCR using FastStart SYBR Green Master Mix and the StepOnePlus™ Real-Time PCR System. To quantify mRNA levels we used a relative quantitation method wherein we constructed a standard curve for each gene using six serial 1:10 dilutions of cDNA pooled from all of our samples. We ran two technical replicates for each standard curve and biological sample and normalized relative Avp and Oxt levels to the housekeeping gene Gapdh after verifying that Gapdh mRNA level did not vary by genotype. The following oligonucleotide primers were used: Avp, TGCGTGTTTCCTGAGCC (forward) and ATGTTGGGTCCGAAGCAG (reverse); Oxt CTGCTACATCCAGAACTGTCC (forward) and AAACAGCCCAGCTCATCG (reverse); Gapdh, GGTAAAGTCATCCCAGAGCTG (forward) and CCTGCTTCACCACCTTCTTG (reverse). Before the RT-qPCR reaction we sequenced the Oxt and Avp PCR products to confirm the specificity of our primers.

#### Data analyses and statistics

The number of animals used was based on previous studies in the field by our group and others, combined with a power analysis. Assumptions of independence, homogeneity of variance, and normality were considered with multiple tests. The assumption of homogeneity of variance was considered through visual inspection of box plots and plots of residuals, and were further tested for homogeneity of variance using Bartlett’s test of homogeneity of variances. Partner preference behaviors in all three mutant backgrounds were considered with paired two-tailed T-test for parametric data or Wilcoxon signed-rank test for non-parametric. Differential time between partner and stranger was calculated as partner huddle duration – stranger huddle duration. The level of statistical significance for each test was set at p = .05. Outliers were detected using a combination of z-score (+/- 3) and plots of original and log-transformed data. One WT OxtR^5^ male (duration with stranger = 3287 seconds), one WT OxtR^1^ male (duration with stranger = 6928 seconds) and one WT OxtR^1^ female (duration with stranger = 7356 seconds) were excluded as outliers due to durations in partner preference assay. Analyses were completed in R (version 1.1.423) and GraphPad Prism (version 7.03).

