## Supplemental Figures for "Oxytocin receptor is not required for social attachment in prairie voles"

### Figure S1

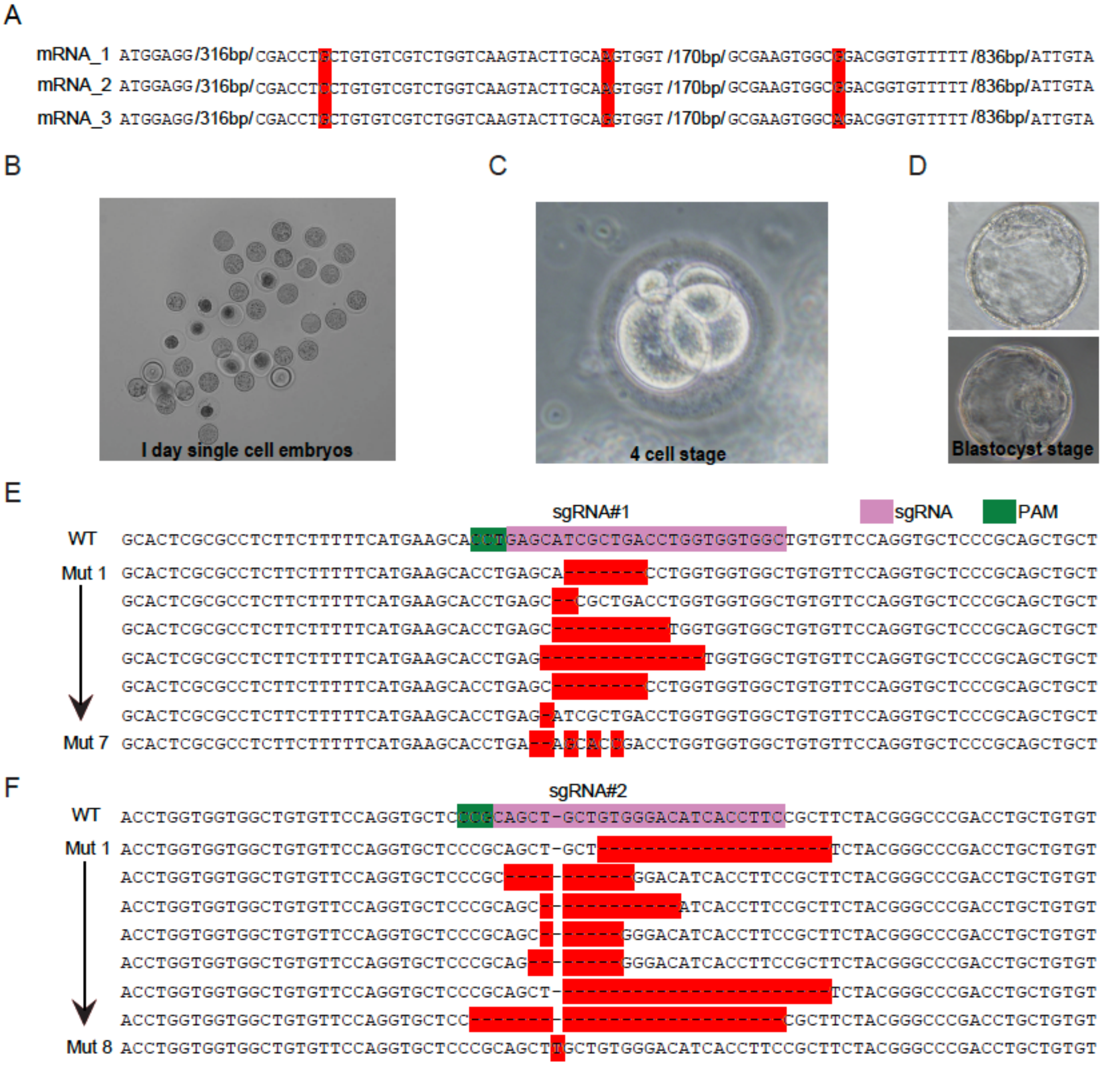


**Figure S1. CRISPR mutagenesis and culture of vole embryos, related to Figure 1**

**A.** 3 naturally occurring synonymous substitutions in *Oxtr* exon1 from 4 wildtype voles.

**B.** Embryos harvested from female voles prior to pronuclear injection.

**C.** Embryo at the 4 cell stage 1 day after ribonucleoprotein complex injection.

**D.** Blastocysts developed in vitro 4 days after injection.

**E.** Sequences from individual blastocysts after injection with sg#1.

**F.** Sequences from individual blastocysts after injection with sg#2.

### Figure S2

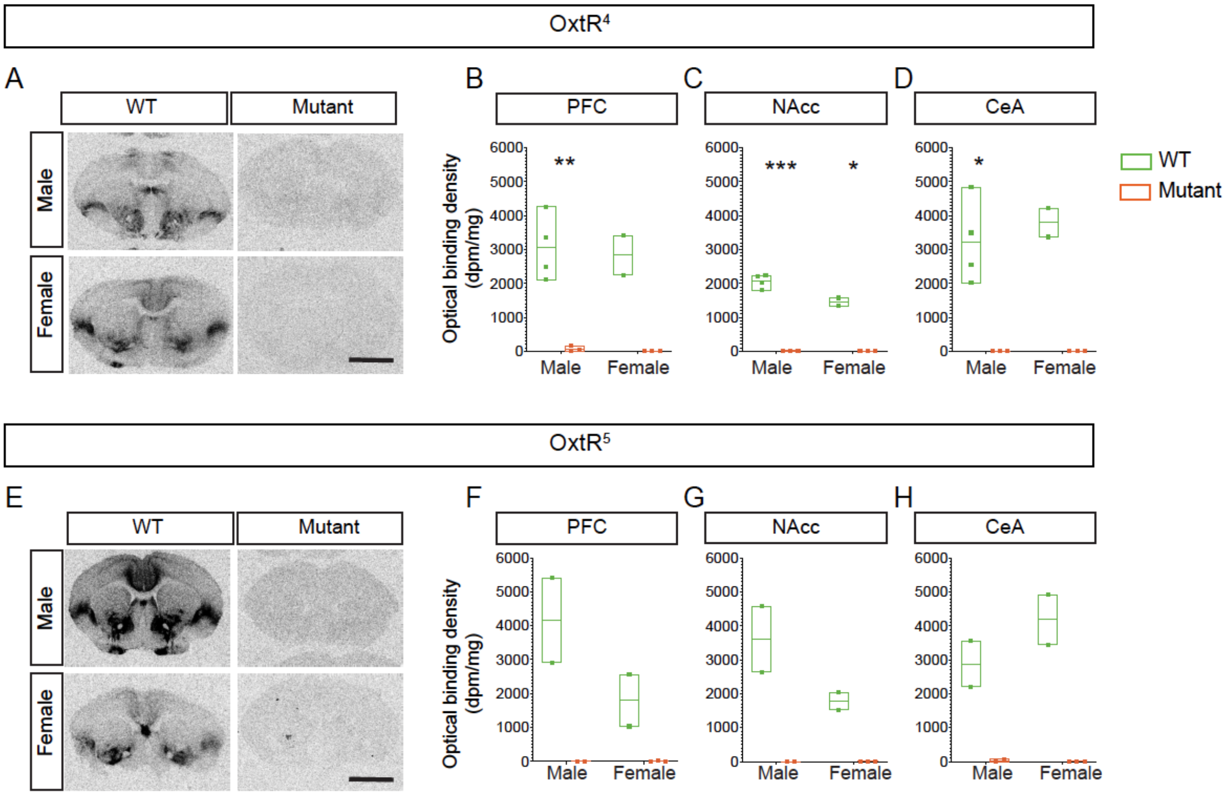


**Figure S2. OxtR4 and OxtR5 products do not bind ligand, related to Figure 2**

**A,E.** Loss of binding with the competitive agonist $^{125}$I-OVTA visualized in coronal sections through the rostral telencephalon of OxtR$^{4–/–}$ (A) and OxtR$^{5–/–}$ (E) voles.

**B-D, F-H.** Optical density based quantification of binding to $^{125}$I-OVTA shows that binding in OxtR$^{4–/–}$ and OxtR$^{5–/–}$ voles is essentially undetectable in PFC (B ,F), NAcc (C, G), and CeA (D, H). Max-Min, Midline denotes mean; n = 4 WT and 3 mutant males, 2 WT and 3 mutant females (B, C, D), 2 WT and mutant males each, 2 WT and 3 mutant females (F, G, H); *$p<0.05$, **$p<0.01$, ***$p<0.001$.

**Figure S3**

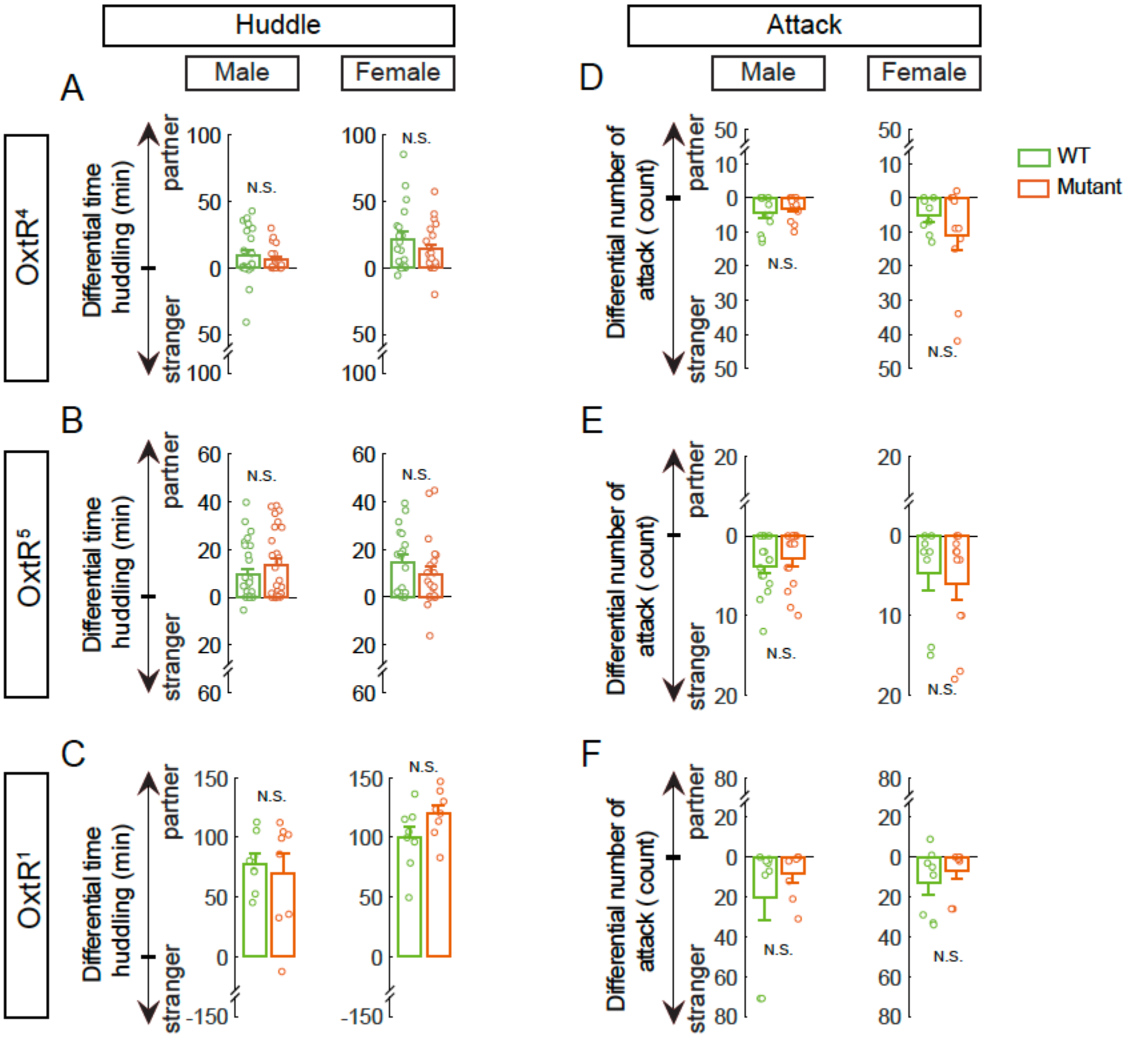


**Figure S3. Oxtr null voles form pair bonds similar to their WT siblings, related to Figure 3**

**A-C.** No difference in time spent huddling differentially with partner over stranger of opposite sex between WT voles and $OxtR^{4-/-}$ (A), $OxtR^{5-/-}$ (B), or $OxtR^{1-/-}$ (C) voles. For individual experimental voles, differential time huddling = duration huddling with partner – duration huddling with stranger.

**D-F.** No difference in number spent differentially attacking stranger of opposite sex between WT, $OxtR^{4-/-}$ (D), $OxtR^{5-/-}$ (E) and $OxtR^{1-/-}$ (F) voles. For individual experimental voles, differential number attacking = number attacking partner – number attacking stranger.

Mean ± SEM; n = 25 WT and 21 mutant males, 19 WT and 21 mutant females (A), 27 WT and mutant males each,19 WT and 21 mutant females (B), 7 WT and 8 mutant males, 7 WT and 8 mutant females each (C), 12 WT and mutant males each, 8 WT and 11 mutant females (D), 15WT and mutant males each, 8 WT and 11 mutant females (E); 8WT and mutant males and females each (F); *p<0.05, **p<0.01, ***p<0.001; N.S., not significant.

**Figure S4**

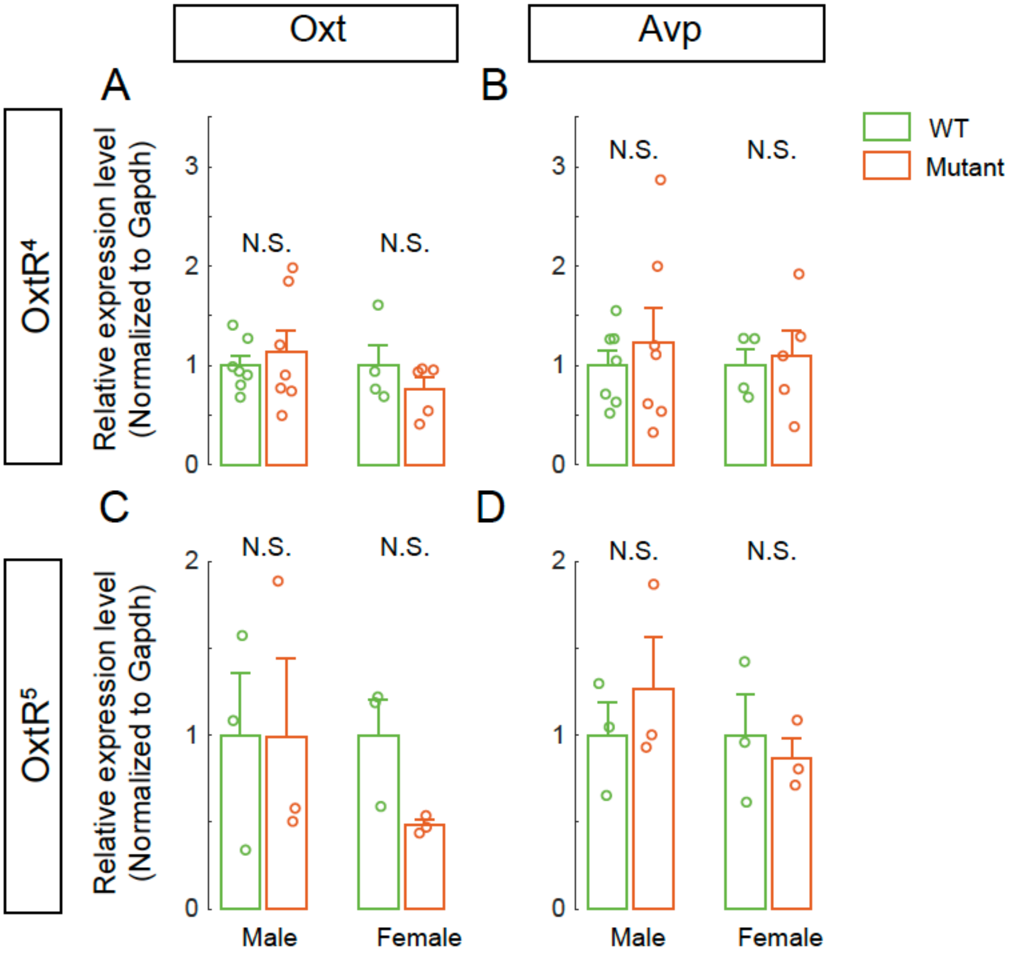


**Figure S4. No increase in oxytocin or AVP levels in Oxtr null voles, related to Figure 4**

**A, B.** No compensatory increase in Oxytocin (A) or AVP (B) mRNA levels in $OxtR^{4-/-}$ voles.

**C, D.** No compensatory increase in Oxytocin (C) or AVP (D) mRNA levels in $OxtR^{5-/-}$ voles.

Mean ± SEM; n = 7 WT and mutant males each, 4 WT and 5 mutant females (A), 7 WT and 6 mutant males, 4 WT and 5 mutant females (B), 8 WT and mutant males each, 9 WT and mutant females each (C-D); N.S., not significant.
